# Immune gene expression changes more during a malaria transmission season than between consecutive seasons

**DOI:** 10.1101/2024.03.20.585963

**Authors:** Kieran Tebben, Salif Yirampo, Drissa Coulibaly, Abdoulaye K. Koné, Matthew B. Laurens, Emily M. Stucke, Ahmadou Dembélé, Youssouf Tolo, Karim Traoré, Amadou Niangaly, Andrea A. Berry, Bourema Kouriba, Christopher V. Plowe, Ogobara K Doumbo, Kirsten E. Lyke, Shannon Takala-Harrison, Mahamadou A. Thera, Mark A. Travassos, David Serre

## Abstract

*Plasmodium* parasites caused over 600,000 deaths in 2022. In Mali, *P. falciparum* is responsible for the majority of malaria cases and deaths and is transmitted seasonally. Anti-malarial immunity develops slowly over repeated exposures to *P. falciparum* but some aspects of this immunity (e.g., antibody titers) wane during the non-transmission, dry season. Here, we sequenced RNA from 33 pediatric blood samples collected during *P. falciparum* infections at the beginning or end of a transmission season and characterized the host and parasite gene expression profiles of paired, consecutive infections. Our analyses showed that human gene expression changes more over the course of one transmission season than it does between seasons, with signatures consistent with the partial development of adaptive immunity during one transmission season, contrasting with the stability in gene expression during the dry season. By contrast, *P. falciparum* gene expression did not seem to vary significantly and remained stable both across and between seasons. Overall, our results provide novel insights into the dynamics of anti-malarial immunity development over short timeframes.

## Introduction

In 2022, malaria caused over 600,000 deaths worldwide^1^. This mortality is primarily caused by *Plasmodium falciparum* infections in children under five years old^2^, who lack protective immunity. Repeated exposure to malaria leads, first to development of immunity to severe malaria (typically occurring in early childhood) and, in later childhood, to immunity against clinical symptoms altogether^3, 4^. Prior studies have demonstrated development of both a cellular^5^ and humoral^6^ response to malaria upon repeated exposures. Memory CD4+ T cells specific for *Plasmodium* blood-stage antigens and skewed towards several T cell phenotypes (e.g., Th1, Tfh, Treg) have been observed^7, 8^, but their role in protective antimalarial immunity remains controversial. In mouse models, Th1 cytokine-biased memory cells appear to protect against malaria^9^, but further work is needed to characterize human T cell memory-mediated protection. Memory B cell populations specific for blood-stage antigens have also been shown to develop with age and exposure^10^ in a transmission-dependent pattern^11^, leading to secretion of *Plasmodium-*specific antibodies^12–15^ that contribute to controlling the parasitemia.

However, the development of the adaptive immune memory response may be complicated by inefficient priming of T cells by antigen presenting cells^4^, dampening of the immune response by regulatory T cells^16^, dysregulation of B and T cells^4, 17^ or development of atypical memory B cell phenotypes^18–20^. In addition, anti-malarial immunity wanes during periods of low exposure^21–24^, but the time scale of this waning and its underlying mechanisms remain unclear^25^.

While individual acquisition and loss of anti-malarial immunity has been studied longitudinally over years, the kinetics of development and loss of anti-malarial immunity over both long- and short-time frames are still incompletely understood. Additionally, the parasite response to changing immune pressure in an infected human during these short periods remains elusive. Previous work has characterized the immune gene expression changes^26^ associated with high and low numbers of repeated clinical malaria episodes across an eight year period^27^, while changes in the expression of *P. falciparum* variant surface antigen, PfEMP1, have been linked to changes in immune status^28, 29^. Since the *P. falciparum* blood-stages are responsible for all clinical symptoms of malaria and since we have access to peripheral blood to examine the human immune response at this stage, studying host and parasite gene expression from infected blood can provide information on how peripheral malaria immunity develops over one transmission season, whether this immunity wanes during the dry season, and how the parasite responds to these changes.

In Bandiagara, Mali, malaria transmission is intensely seasonal, with a high transmission wet season from June to December and a low transmission dry season from January to May. Each child 0-14 years of age experiences on average 2.2 clinical malaria episodes during the high transmission season, compared to 0.275 during the low transmission season^30^. This high seasonality makes Bandiagara an ideal location to study the dynamics of antimalarial immunity development and loss over short time frames. Here, we use dual RNA-sequencing analyses of whole blood samples collected during symptomatic *P. falciparum* infections that occurred i) at the beginning and end of one transmission season and ii) at the end and beginning of two consecutive transmission seasons, to study the dynamics of the anti-malarial immune response over a short time scale.

## Results

### Dual RNA-sequencing to characterize human and P. falciparum gene expression

We extracted and sequenced RNA from whole blood samples collected during 33 symptomatic *P. falciparum* infections from 11 Malian children, aged 1-10 years (Table 1). All samples were collected during a patient-initiated visit due to self-identified malaria symptoms (e.g., fever, headache) and for which *Plasmodium* parasitemia was confirmed by light microscopy^30^ (**Table 1**). The mean parasitemia was 64,338 parasites per µL of blood (range 225 – 198,325).

**Table 1:** Sample characteristics of the selected participants.

| Early vs. Late analysis group |  |  |  |  |  |  |  |  |  |
| --- | --- | --- | --- | --- | --- | --- | --- | --- | --- |
| Participant ID | Sex | Ethnicity | Age* (years) | Collection Date (dd-mm-yyyy) | Collection Season | Parasitemia (parasites per μL) | Temp. ($^{\circ}$C) | Hgb conc. (g/dL) | Complexity of infection |
| A | F | Dogon | 7 | 20-Aug-2010 | Early wet | 36,375 | 39.5 | 10.6 | Monoclonal |
|  |  |  |  | 02-Nov-2010 | Late wet | 225 | 39.6 | 10.7 | Monoclonal |
| B | M | Dogon | 7 | 18-Sept-2009 | Early wet | 56,600 | 37.5 | 11.3 | Polyclonal |
|  |  |  |  | 17-Sept-2010 | Early wet | 83,075 | 36.1 | 10.6 | Polyclonal |
|  |  |  |  | 17-Nov-2009 | Late wet | 97,125 | 37.5 | 10.3 | Monoclonal |
|  |  |  |  | 21-Nov-2010 | Late wet | 12,450 | 37.9 | 9.0 | Polyclonal |
| C | F | Dogon | 2 | 29-Sept-2010 | Early wet | 179,550 | 39.8 | 7.7 | Monoclonal |
|  |  |  |  | 27-Nov-2010 | Late wet | 65,100 | 38.9 | 11.6 | Monoclonal |
| D | M | Dogon | 5 | 13-Aug-2012 | Early wet | 181,800 | 38.2 | 11.2 | Polyclonal |
|  |  |  |  | 09-Nov-2012 | Late wet | 39,975 | 38.4 | 11.6 | Polyclonal |
| E | M | Dogon | 5 | 29-Sept-2010 | Early wet | 118,500 | 37.7 | 11.1 | Monoclonal |
|  |  |  |  | 25-Dec-2010 | Late wet | 120,400 | 38.4 | 10.5 | Polyclonal |
| F | M | Dogon | 3 | 03-Sept-2010 | Early wet | 37,206 | 38.4 | 9.7 | Monoclonal |
|  |  |  |  | 20-Nov-2010 | Late wet | 2,100 | 38.6 | 6.6 | Monoclonal |
| G | M | Peuhl | 5 | 17-Sept-2010 | Early wet | 91,350 | 38.8 | 12.1 | Polyclonal |
|  |  |  |  | 12-Dec-2010 | Late wet | 48,325 | 39.3 | 11.2 | Polyclonal |

| Late vs. Early analysis group |  |  |  |  |  |  |  |  |  |
| --- | --- | --- | --- | --- | --- | --- | --- | --- | --- |
| Participa<br>nt ID | Se<br>x | Ethnici<br>ty | Age*<br>(year<br>s) | Collection<br>Date (dd-mm-yyyy) | Collectio<br>n<br>Season | Parasitemia<br>(parasites per<br>μL) | Temp.<br>(°C) | Hgb conc<br>.<br>(g/dL<br>) | Complexi<br>ty of<br>infection |
| A | F | Dogon | 6 | 20-Aug-2010 | Early<br>wet | 36,375 | 39.5 | 10.6 | Monoclon<br>al |
|  |  |  |  | 12-Dec-2009 | Late wet | 900 | 39.8 | 9.9 | Polyclona<br>l |
| B | M | Dogon | 6 | 17-Nov-2009 | Late wet | 97,125 | 37.5 | 10.3 | Monoclon<br>al |
|  |  |  |  | 17-Sept-2010 | Early<br>wet | 83,075 | 36.1 | 10.6 | Polyclona<br>l |
| C | F | Dogon | 1 | 19-Dec-2009 | Late wet | 10,800 | 35.9 | 9.0 | Monoclon<br>al |
|  |  |  |  | 29-Sept-2010 | Early<br>wet | 179,550 | 39.8 | 7.7 | Polyclona<br>l |
|  |  |  |  | 27-Nov-2010 | Late wet | 65,100 | 38.9 | 11.6 | Monoclon<br>al |
|  |  |  |  | 12-Aug-2011 | Early<br>wet | 71,900 | 38.1 | 11.7 | Polyclona<br>l |
| E | M | Dogon | 5 | 25-Dec-2010 | Late wet | 120,400 | 38.4 | 10.5 | Polyclona<br>l |
|  |  |  |  | 26-Jul-2011 | Early<br>wet | 198,325 | 39.1 | 12.2 | Polyclona<br>l |
| G | M | Peuhl | 4 | 18-Dec-2009 | Late wet | 1,125 | 38.3 | 9.7 | Monoclon<br>al |
|  |  |  |  | 17-Sept-2010 | Early<br>wet | 91,350 | 38.8 | 12.1 | Polyclona<br>l |
| H | M | Dogon | 10 | 26-Nov-2010 | Late wet | 47,475 | 36.2 | 10.9 | Monoclon<br>al |
|  |  |  |  | 11-Sept-2011 | Early<br>wet | 30,475 | 37.5 | 10.8 | Polyclona<br>l |
| I | M | Dogon | 2 | 21-Dec-2010 | Late wet | 186,350 | 38.3 | 10.8 | Polyclona<br>l |
|  |  |  |  | 26-Sept-2011 | Early<br>wet | 43,800 | 38.9 | 10.3 | Monoclon<br>al |
| J | M | Dogon | 2 | 1-Nov-2009 | Late wet | 43,800 | 39.4 | 8.5 | Polyclona<br>l |
|  |  |  |  | 20-Aug-2010 | Early<br>wet | 37,550 | 37.6 | 6.4 | Monoclon<br>al |
| K | M | Dogon | 2 | 23-Dec-2010 | Late wet | 15,100 | 40 | 9.8 | Polyclona<br>l |
|  |  |  |  | 30-Sept-2011 | Early<br>wet | 62,100 | 39.7 | 11.9 | Monoclon<br>al |
| *age at first sampled infection. |  |  |  |  |  |  |  |  |  |

We included in this analysis blood samples from children that had 1) two *P. falciparum* symptomatic infections in the same transmission season, one at the beginning of the wet season and one at the end (n=8 pairs) (Early vs. Late comparison, in blue on **Figure 1**), and/or 2) two *P. falciparum* symptomatic infections in consecutive years, one at the end of the transmission season of year 1 and one at the beginning of the transmission season of year 2 (n=11 pairs) (Late vs. Early comparison, in red on **Figure 1**) (**Table 1**).

**Figure 1:**
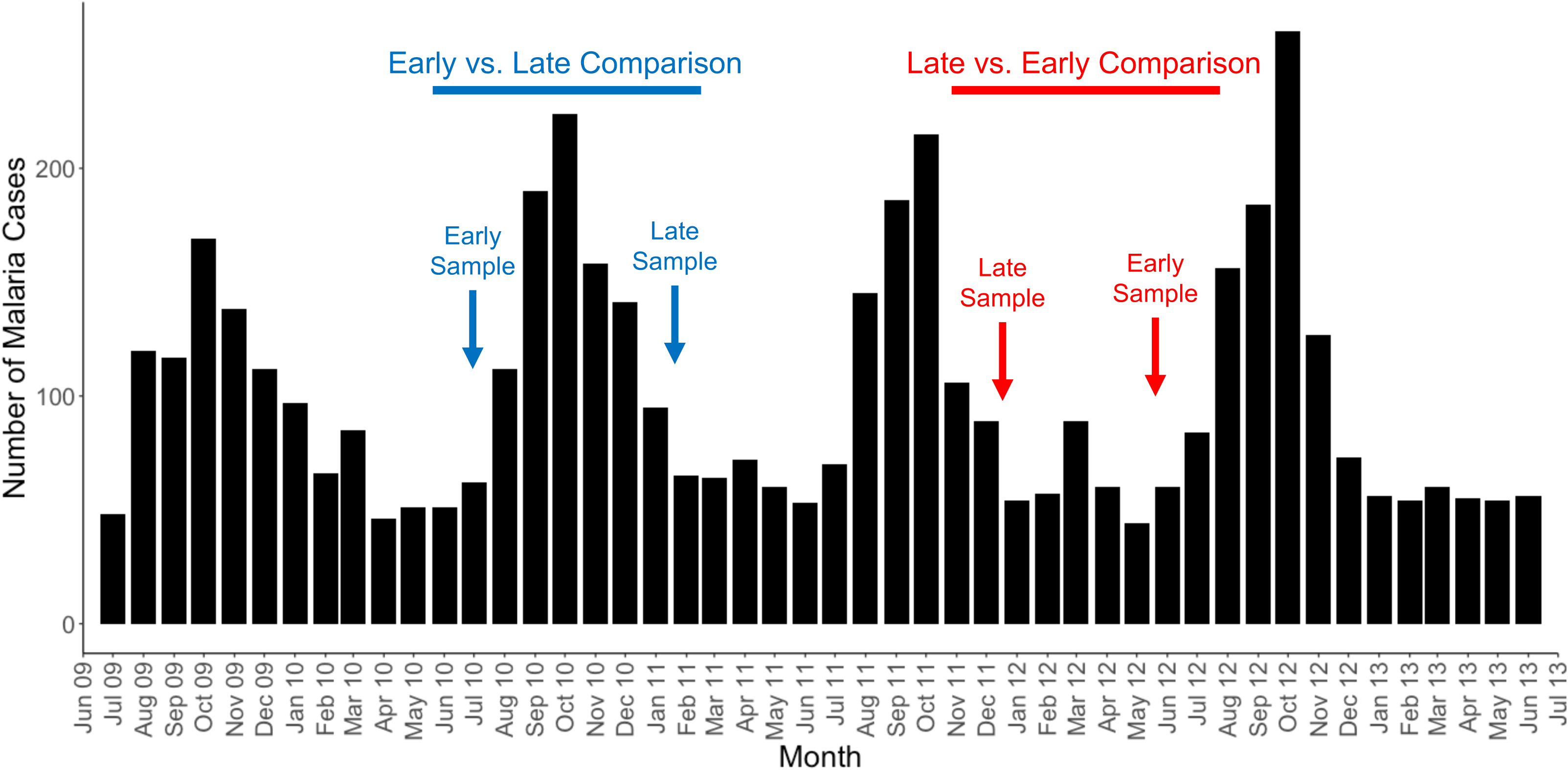
Schematic of the sampling strategy. The black bars show the number of symptomatic malaria cases reported in the entire longitudinal cohort^30^ across four years. The blue and red arrows illustrate the sampling strategy of paired infections (i.e., from the same child) that would be selected for, respectively, Early vs. Late comparisons (to examine the development of immunity over one transmission season) and Late vs. Early comparisons (to examine the loss of immunity over one dry season).

To confirm that *P. falciparum* caused all infections, we first mapped all reads to the genomes of *P. falciparum, P. vivax, P. ovale and P. malariae,* simultaneously, and found that more than 98% of *Plasmodium* reads mapped to the *P. falciparum* genome in each sample (**Supplemental Table 1**). We then mapped all reads to the human and *P. falciparum* genomes, simultaneously. We obtained 32-139 million reads mapping to human (49% to 99%) and 0.3 to 50 million reads mapping to *P. falciparum* (0.3% to 50%), allowing robust characterization of host and parasite transcriptomes **(Supplemental Table 1**).

### Late season symptomatic infections are characterized by a stronger adaptive immune response

We first compared the human gene expression profiles generated from symptomatic infections, from the same child, at the beginning and at the end of one transmission season to investigate potential differences in immune response (n=8 pairs, **Table 1**). Of 9,181 expressed human genes, 130 genes were significantly differentially expressed (FDR<0.1) between symptomatic infections occurring early versus late in the season, after adjusting for parasitemia (**Supplemental Figure 1A, Supplemental Table 2**). Interestingly, genes with functions indicative of an adaptive immune response, such as T cell activation (e.g., CCL5^31^, ADA^32^) and T and NK cell granules (e.g., GNLY^33^, FGFBP2^34^), were significantly increased in expression during late season infections. By contrast, genes with functions indicative of an innate immune response, such as pro-inflammatory cytokines (e.g., IL-18^35^), interferon-stimulated genes (e.g., GBP1^36^, GBP4^36^, GBP5^36^, PARP14^37^) and regulators of the innate immune system (e.g., CLIC4^38^, LRRK2^39^), were significantly decreased in expression during late season infections.

Overall, this result is consistent with partial development of adaptive immunity to parasites over repeated exposures throughout the season.

To determine whether these differences in gene expression resulted from changes in white blood cell proportion or true differences in gene regulation, we estimated the relative proportion of each immune cell type using gene expression deconvolution^40^ and adjusted our differential expression analyses for the proportion each cell type (**Supplemental Table 2**). After adjusting for cell composition, only one gene, myosin light chain 9 (MYL9), remained differentially expressed between early and late season infections, with a higher expression in late season infections (**Supplemental Figure 1B, Supplemental Table 2**). As a myosin molecule, MYL9 has diverse roles in different cell types, and can interact with the T cell activation marker CD69 to induce inflammation during infections^41^. MYL9 has been reported by one study to be expressed during treatment and recovery from malaria^42^, and could potentially be involved in promoting an adaptive immune response to infection.

We then examined which immune cell types differed in relative proportion between early and late season symptomatic infections in the same individual. On average, we found that late season infections were characterized by a higher proportion of adaptive immune cells, whereas early season infections were characterized by a higher proportion of innate immune cells.

Specifically, late season infections had proportionally more naïve B cells, CD8 T cells, and resting NK cells than early season infections (p < 0.03, **Figure 2A**). In contrast, early season infections had proportionally more activated NK cells, neutrophils, resting mast cells, plasma cells, and activated dendritic cells (p < 0.05, **Figure 2B**). These observations are consistent with a greater role of an innate response in early season infections, while the adaptive immune response dominates late in the transmission season, suggesting that there is a significant acquisition of anti-malarial immunity, even on this short time scale.

**Figure 2:**
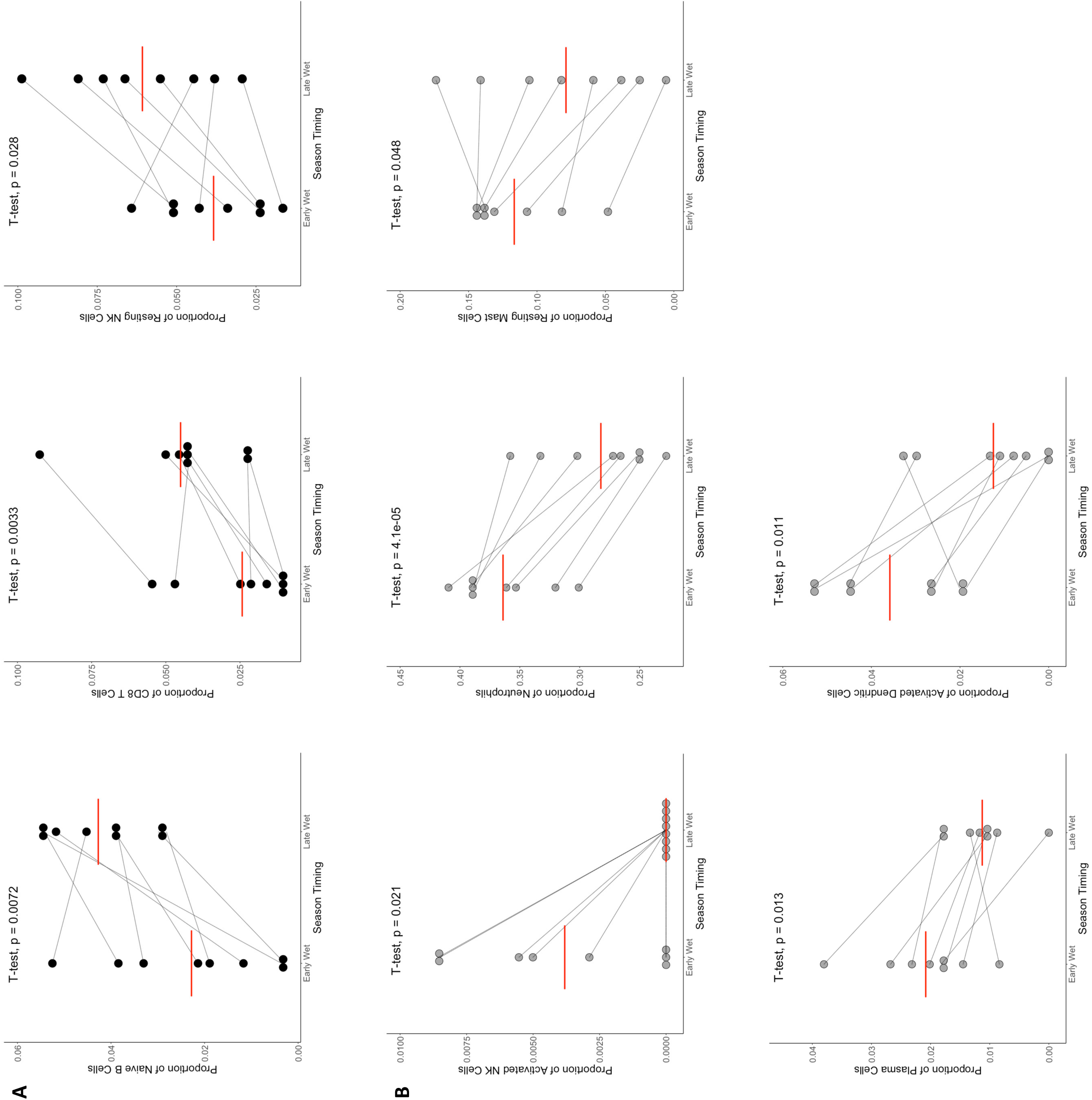
Change in the relative proportion of immune cell types between symptomatic infections occurring early and late in the transmission season. Each panel shows the proportion of one WBC subset estimated by gene expression deconvolution, with the thin black lines joining estimates from the same individual. **A) Cell types that are enriched in late season infections.** The panels correspond, from left to right, to naïve B cells, CD8 T cells, resting NK cells. **B) Cell types that are enriched in early season infections.** The panels correspond, from left to right and top to bottom, to activated NK cells, neutrophils, resting mast cells, plasma cells and activated dendritic cells. All comparisons utilize student paired T-test with significance defined as p ≥ 0.05. Note the difference in y-axis scale due to differences in the proportion of each immune cell subtype.

### Changes in human gene expression are minimal across the dry season

To begin to understand whether the gene expression associated with developing immunity changes during the dry season (i.e., between transmission seasons), we compared gene expression profiles of 11 pairs of samples from the same children collected during one symptomatic infection at the end of one transmission season and during one symptomatic infection at the beginning of the next transmission season (**Figure 1**, **Table 1**, **Supplemental Table 2**). In contrast to gene expression changes observed between infections occurring in the beginning and end of the same season, and despite the larger sample size (11 vs. 8), we only identified one gene (MARCO, a macrophage receptor) whose expression was significantly different in this comparison (**Supplemental Figure 2**). This suggests that immunity remains relatively stable during the dry season and that the immune gene expression in response to parasites in the next season is very similar to that of the end of the previous season, in the absence of interval exposure to infected mosquitoes.

Despite the lack of detectable gene expression differences, we analyzed how proportions of immune cells may have changed between transmission seasons (**Supplemental Table 2**). We found that infections occurring late in one transmission season had significantly more naïve B cells than infections occurring early in the subsequent transmission season (**Figure 3**). This supports our above findings that late season symptomatic infections are characterized by a more adaptive immune cell signature, but the overall immune response is stable across the dry season. These data suggest that there is no detectable waning of immunity over the dry season, at least as measured by gene expression among immune cells detectable in the peripheral blood.

**Figure 3:**
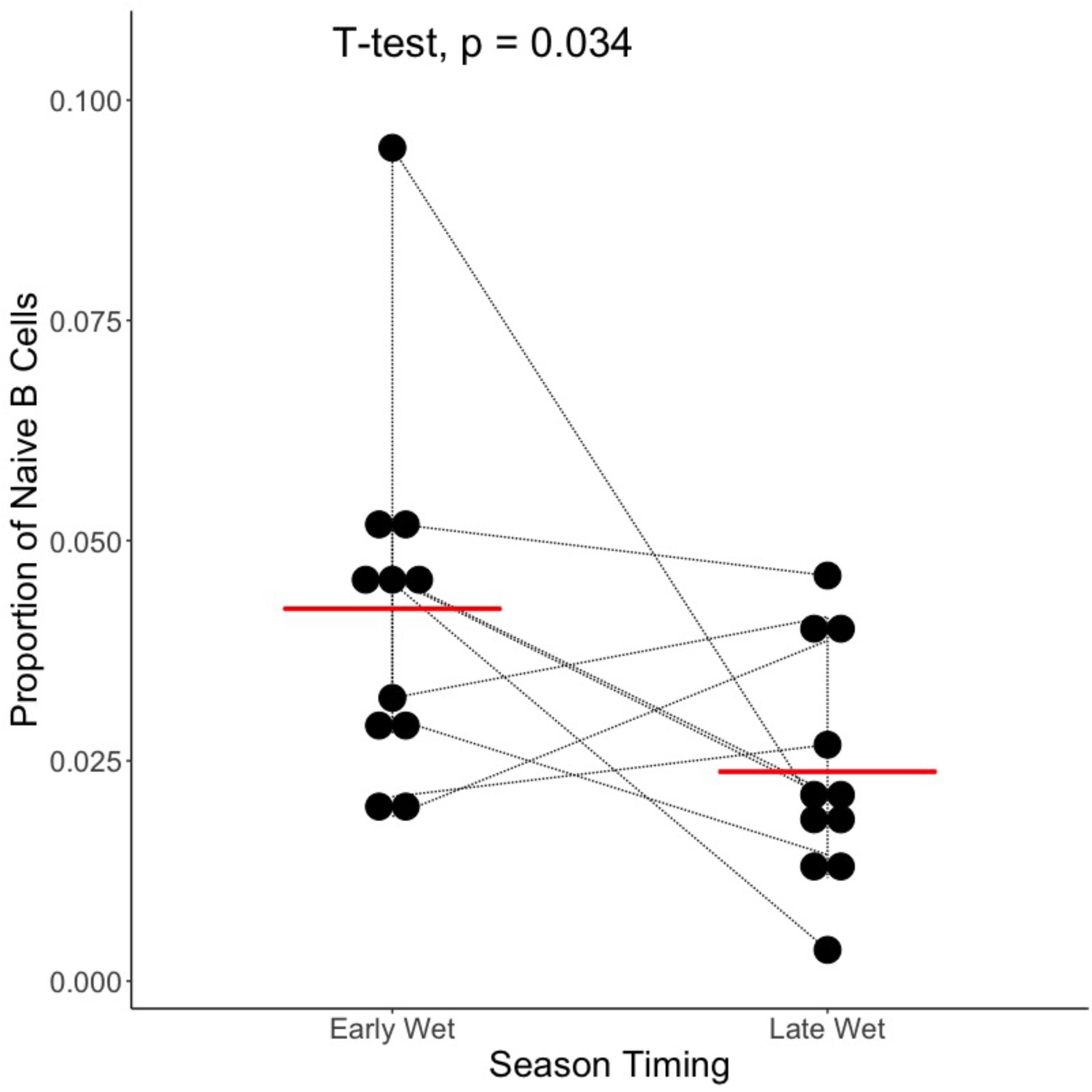
Change in the relative proportion of naïve B cells between symptomatic infections occurring late and early in subsequent transmission seasons. The panel shows the proportion of naïve B cells estimated by gene expression deconvolution, with the thin black lines joining estimates from the same individual, compared with a paired T-test.

### P. falciparum gene expression varies minimally over the course of a transmission season or between transmission seasons

Since circulating *P. falciparum* parasites are exposed to the immune system in the blood, we might expect that, as the immune response changes over a transmission season, parasites would vary their gene expression to adapt to changing immune pressures. We first compared the expression of 2,574 *P. falciparum* genes from the 8 pairs of samples selected from the beginning and end of one transmission season (**Table 1**). Interestingly, compared to more than 100 differentially expressed human genes between these samples, we identified only nine parasite genes whose expression differed after adjustment for parasitemia (**Supplemental Table 3, Supplemental Figure 3**). This suggests that, despite changes in host immunity, parasite transcriptional programs in peripheral circulation remain very similar, but may not reflect that of liver-resident parasites in situ. Of these nine differentially expressed genes, only three had annotated functions and two of the genes, PfPTP3 and PfPTP7, are involved in trafficking the variant surface antigen PfEMP1 to the RBC surface^43, 44^ (the third one, PfFRM2, is involved in daughter merozoite formation ^45^). This observation is interesting since *P. falciparum* has been shown, *in vitro*, to vary PfEMP1 expression in response to environmental changes^29^, which likely includes host immune status. Additionally, one study of Kenyan children identified particular PfEMP1 subtypes associated with immune status to severe malaria^28^. An important limitation of the current study is that we did not analyze expression of the different PfEMP1 genes (due to the high sequence homology between PfEMP1 genes and high variability among parasites, it is difficult to map and analyze rigorously PfEMP1s from short read RNA-seq data). It will be important to follow up on this observation and characterize in future studies whether particular PfEMP1 subtypes are expressed at different times during the transmission season or at different levels of host immunity.

Because of potential changes in host immune pressure over the course of the dry season, we also compared parasite gene expression from 11 pairs of samples selected from the end of one transmission season and the beginning of the next transmission season. Of the 2,483 parasite genes expressed, we did not detect any differentially expressed genes in this analysis (**Supplemental Table 3, Supplemental Figure 4**). This could support our above results that the host immune response is stable, in the absence of ongoing *P. falciparum* exposure, between transmission seasons and *P. falciparum* parasites are exposed to similar environments during clinical infections. (We also did not observe any differences in the developmental stage composition between early and late samples from the same season, nor between late samples from one season and early samples from the next (**Supplemental Table 3**).

## Discussion

Overall, our data suggest that adaptive immunity to *P. falciparum* partially develops over the course of one transmission season, with evidence from gene expression of activated T and NK cells in late season infections, while we did not detect any waning during the non-transmission (dry) season. Interestingly, despite this change in host immune pressure across the transmission season, we did not detect any substantial changes in the *P. falciparum* gene expression.

Prior work has described the development of adaptive immunity to *P. falciparum* over long time frames^4, 6, 46–48^ and subsequent waning during periods without consistent parasite exposure^21, 22,49^. We observed an increase in naïve B cells, NK cells and CD8 T cells in late season infections, and an adaptive immune gene expression signature, suggesting that some adaptive immunity incrementally develops even within one transmission season. While the memory B cell population has been shown to slowly develop over multiple infections^10^, our observations suggest that the B cell response to *P. falciparum* may begin to develop even over a few exposures (but see some limitations below). NK cells can also produce a memory-like response^50^ and mediate efficient killing of infected RBCs in an adaptive-like response^51, 52^ in cooperation with *P. falciparum*-specific antibodies developed as part of the humoral response^13, 15, 53, 54^. CD8 T cells are implicated in immunity to the liver stages of *P. falciparum*^55^ and their enrichment in late season infections is consistent with developing immunity to this stage throughout the transmission season.

Surprisingly, we did not detect enrichment for memory B and T cells, specifically, which would have suggested anti-malarial immune memory development during one transmission season. This could partially be due to the limited resolution of our gene expression deconvolution technique in distinguishing precisely between memory and naïve lymphocyte populations, the atypical, exhausted phenotype of memory lymphocytes that develop after malaria infection and have unique gene expression profiles^17, 18, 20, 56, 57^, or a true defect in memory cell generation over few infections. Future work with more high-resolution techniques such as flow cytometry will be necessary to confirm our results and disentangle these possibilities. Taken together, our findings could suggest that despite development of an appropriate adaptive response after malaria exposures (i.e., accumulation of B and T cells late in a season), impairment of the memory response to *P. falciparum* potentially occurs even over a few symptomatic infections during one season. Prior work has also suggested that the precise number of infections (symptomatic or asymptomatic) experienced during a transmission season also strongly influences the development of immunity during that season^58, 59^. Future basic immunology work is warranted to validate these findings and further disentangle the relationship between development of immunity and number of infections per season.

Interestingly, despite previous evidence of waning antibody titers over the dry season^21^, we did not detect any appreciable differences in adaptive immune-related gene expression over the course of one non-transmission season. The lack of differences in gene expression and immune cell composition between late season and subsequent early season infections suggests that the response to infecting parasites remains relatively stable between the end of one season and the beginning of the next, in Mali. While the lack of detectable gene expression differences is not proof of the lack of immunologic differences, it is worth noting here that i) the sample size for this analysis was slightly larger than for the analysis of the changes during one season and ii) that the samples spanning one dry season were collected 7-10 months apart (compared to 2-3 months apart for the samples in the same transmission season, where ongoing exposure to *P. falciparum* is occurring). Interestingly, a previous study has reported that parasites can persist as sub-clinical infections through the dry season in Mali, with reportedly little effect on the host immune response^60^ and it is possible that these asymptomatic infections could help to maintain immunity through the dry season.

One important limitation of our current study is its small sample size, which limits overall study power. Individual differences in baseline gene expression, prior malaria exposure and immunity and number of infections experienced during the malaria season, likely impact gene expression and may confound our analyses but are difficult to control for due to the sample size. Indeed, prior work has identified age and number of exposures as important determinants in the development of immunity^49, 58, 61^. The Peuhl ethnicity has also been associated with genetic protection from malaria^62^ and differences in immune gene expression between Dogon and Peuhl individuals could influence our findings. Here, we used paired analyses, comparing samples from the same individual collected during different time points throughout the season, to limit the influence of these individual variations but future work with larger cohorts, which can better control for these potential confounders, will be essential to confirm and strengthen findings presented here.

Additionally, because our samples were collected when a child presented to clinic with symptoms, all infections included in our analyses were identified and treated at different lengths of time after initial symptom presentation. Specifically, since the adaptive immune response takes several weeks to develop, the interval between infection and diagnosis may influence the gene expression profile. Additionally, treatment of infections could influence the development of a productive adaptive immune response, especially if parasite exposure is very short-lived. We also only included individuals in this study who presented with symptomatic disease, which could introduce an important sampling bias by studying only those individuals who did not yet develop anti-disease immunity to malaria.

## Conclusions

In this work, we described the transcriptional profiles of the host and parasite during uncomplicated *P. falciparum* infections occurring at the beginning and end of consecutive transmission seasons. We found that the human immune response changes more over the course of one transmission season than between transmission seasons, despite a lower power (n=8 vs n=11) and a shorter time frame (2-3 months vs 7-10 months). This observation suggests that the immune response to *P. falciparum* changes over a transmission season to adopt an adaptive immune signature later during the transmission season, while it remains relatively stable between transmission seasons. In contrast, we found that *P. falciparum* gene expression varies minimally over this short time scales. Overall, this study contributes new insights into anti-malarial immunity development over repeated exposures during the short time scale of one transmission season. These findings have important implications for understanding the development of protective immunity to malaria that could be exploited by future vaccine and prevention efforts.

## Materials and Methods

### Ethics approval and consent

Individual informed consent/assent was obtained from all children and their parents. The study protocol and consent/assent processes were approved by the institutional review boards of the Faculty of Medicine, Pharmacy and Dentistry of the University of Maryland, Baltimore and of the University of Sciences, Techniques and Technologies of Bamako, Mali (IRB numbers HCR-HP- 00041382 and HP-00085882).

### Samples

We selected 55 whole blood samples, collected directly in PAXgene blood RNA tubes, from children experiencing a symptomatic uncomplicated malaria episode caused by *Plasmodium falciparum* parasites at the beginning or end of the transmission season in Mali (i.e., June to December). The presence of parasites and the parasite species were initially determined by light microscopy using thick blood smears. All infections were successfully treated with antimalarial drugs according to the Mali National Malaria Control Programme standards.

### Case Definition

Children were classified, by the field clinicians, as experiencing symptomatic uncomplicated malaria if they i) sought treatment from the study clinic, ii) experienced symptoms consistent with malaria (i.e., fever, headache, joint pain, abdominal pain, vomiting or diarrhea), and iii) *Plasmodium* parasites were detected, at any density, by thick blood smear, and if they lacked any signs of severe malaria (e.g., coma, seizures, severe anemia) ^30^.

### Generation of RNA-seq data

We extracted RNA from whole blood using MagMax blood RNA kits (Themo Fisher). Total RNA was subjected to rRNA depletion and polyA selection (NEB) before preparation of stranded libraries using the NEBNext Ultra II Directional RNA Library Prep Kit (NEB). cDNA libraries were sequenced on an Illumina NovaSeq 6000 to generate ∼60-156 million paired-end reads of 75 bp per sample. To confirm that *P. falciparum* was responsible for each malaria episode, we first aligned all reads from each sample using hisat2 v2.1.0^63^ to a fasta file containing the genomes of all *Plasmodium* species endemic in Mali downloaded from PlasmoDB ^64^ v55: *P. falciparum* 3D7*, P. vivax* PvP01*, P. malariae* UG01, and *P. ovale curtisi* GH01. After ruling out coinfections and misidentification of parasites, we aligned all reads using hisat2 to a fasta file containing the *P. falciparum* 3D7 and human hg38 genomes i) using default parameters and ii) using (--max-intronlen 5000). Reads mapping uniquely to the hg38 genome were selected from the BAM files generated with the default parameters. Reads mapping uniquely to the *P. falciparum* genome were selected from the BAM files generated with a maximum intron length of 5,000 bp. PCR duplicates were removed from all files using custom scripts. We then calculated read counts per gene using gene annotations downloaded from PlasmoDB (*P. falciparum* genes) and NCBI (human genes) and the subread featureCounts v1.6.4^65^.

### Gene expression analysis

Read counts per gene were normalized into counts per million (CPM), separately for human and *P. falciparum* genes. Only human or *P. falciparum* genes that were expressed at least at 10 CPM in > 50% of the samples were retained for further analyses. Read counts were normalized via TMM for differential expression analyses. Statistical assessment of differential expression by the time during the season in which a sampled infection occurred was conducted, separately for the human and *P. falciparum* genes, in edgeR (v 3.32.1)^66^ using a quasi-likelihood negative binomial generalized model. We used a paired design to make intra-individual comparisons of gene expression between early and late season infections to minimize interindividual effects such as differing levels of developed immunity due to age and exposure history. All models were corrected for the parasitemia of each infection. Adjusted models were corrected for the cell composition of each sample (see below). All gene expression analyses were corrected for multiple testing using FDR ^67^ (FDR = 0.1).

### Gene expression deconvolution

CIBERSORTx ^40^ was used to estimate, in each sample, the proportion of i) human immune cell types and ii) *Plasmodium* developmental stages. To deconvolute human gene expression profiles, we used as a reference LM22 ^68^, a validated leukocyte gene signature matrix using 547 genes to differentiate 22 immune subtypes. A custom signature matrix derived from *P. berghei* scRNA-seq data was used for *P. falciparum* stage deconvolution, using orthologous genes between the two species ^69^. Relative proportions of each human immune cell type and *P. falciparum* blood stage from each sample are available in **Supplemental Table 4.**

### Statistical Analysis

All statistical analyses not mentioned above were conducted in R (version 4.0.3). Paired t-tests were used to compare cell proportions between groups.

### Complexity of infection

We used samtools^70^ mpileup to call the genotype at each sequenced position in all samples directly from the RNA-seq reads. We removed positions within *Plasmodium* multi-gene families due to inaccurate mapping of reads within these regions because of high sequence variability. We then calculated the reference allele frequency (RAF) at each position directly from the resulting files. To determine the complexity of each infection (i.e., monoclonal vs. polyclonal), we visualized graphically the distribution of RAF in each sample. Samples with a U-shaped curve, with the RAF for most positions being either 0 or 1, were considered monoclonal.

Samples with RAF between 0 and 1, representing a substantial deviation from the U-shaped curve, were considered polyclonal^71^.

## Data and Code Availability

All sequence data generated in this study are deposited in the Sequence Read Archive under the BioProject XXX. Custom scripts are available at https://github.com/tebbenk/seasonality.

## Figure Legends

**Supplemental Figure 1: Differences in host gene expression between infections occurring early and late during one transmission season.** Each point represents one gene plotted according to the fold-change and the p-value. Red points represent genes that are more highly expressed in late season infections. Blue points represent genes that are more highly expressed in early season infections. **A)** Before adjustment for cell composition **B)** After adjustment for cell composition.

**Supplemental Figure 2: Differences in host gene expression between infections occurring late in one transmission season and early in the next, unadjusted for cell composition.** Each point represents one gene plotted according to the fold-change and the p-value. Red points represent genes that are more highly expressed in late season infections. Blue points represent genes that are more highly expressed in early season infections.

**Supplemental Figure 3: Differences in *P. falciparum* gene expression between infections occurring early and late during one transmission season, unadjusted for cell composition.** Each point represents one gene plotted according to the fold-change and the p-value. Red points represent genes that are more highly expressed in late season infections. Blue points represent genes that are more highly expressed in early season infections.

**Supplemental Figure 4: Differences in *P. falciparum* gene expression between infections occurring late during one transmission season and early in the next, unadjusted for cell composition.** Each point represents one gene plotted according to the fold-change and the p-value. Red points represent genes that are more highly expressed in late season infections. Blue points represent genes that are more highly expressed in early season infections.

## References

1. WHO GMP. World Malaria Report 2023. 2023.

2. Phillips MA, Burrows JN, Manyando C, van Huijsduijnen RH, Van Voorhis WC, Wells TNC. Malaria. Nat Rev Dis Primers. 2017;3:17050. Epub 20170803. doi: 10.1038/nrdp.2017.50. PubMed PMID: 28770814.

3. Achtman AH, Bull PC, Stephens R, Langhorne J. Longevity of the Immune Response and Memory to Blood-Stage Malaria Infection. In: Langhorne J, editor. Immunology and Immunopathogenesis of Malaria. Berlin, Heidelberg: Springer Berlin Heidelberg; 2005. p. 71–102.

4. Langhorne J, Ndungu FM, Sponaas AM, Marsh K. Immunity to malaria: more questions than answers. Nat Immunol. 2008;9(7):725–32. doi: 10.1038/ni.f.205. PubMed PMID: 18563083.

5. Good MF, Miller LH. Involvement of T cells in malaria immunity: implications for vaccine development. Vaccine. 1989;7(1):3–9. doi: 10.1016/0264-410x(89)90002-9. PubMed PMID: 2655342.

6. Rogers KJ, Vijay R, Butler NS. Anti-malarial humoral immunity: the long and short of it. Microbes Infect. 2021;23(4-5):104807. Epub 20210305. doi: 10.1016/j.micinf.2021.104807. PubMed PMID: 33684519; PMCID: PMC8292161.

7. Kurup SP, Butler NS, Harty JT. T cell-mediated immunity to malaria. Nat Rev Immunol. 2019;19(7):457–71. doi: 10.1038/s41577-019-0158-z. PubMed PMID: 30940932; PMCID: PMC6599480.

8. Soon MSF, Lee HJ, Engel JA, Straube J, Thomas BS, Pernold CPS, Clarke LS, Laohamonthonkul P, Haldar RN, Williams CG, Lansink LIM, Moreira ML, Bramhall M, Koufariotis LT, Wood S, Chen X, James KR, Lonnberg T, Lane SW, Belz GT, Engwerda CR, Khoury DS, Davenport MP, Svensson V, Teichmann SA, Haque A. Transcriptome dynamics of CD4(+) T cells during malaria maps gradual transit from effector to memory. Nat Immunol. 2020;21(12):1597–610. Epub 20201012. doi: 10.1038/s41590-020-0800-8. PubMed PMID: 33046889.

9. Opata MM, Ibitokou SA, Carpio VH, Marshall KM, Dillon BE, Carl JC, Wilson KD, Arcari CM, Stephens R. Protection by and maintenance of CD4 effector memory and effector T cell subsets in persistent malaria infection. PLoS Pathog. 2018;14(4):e1006960. Epub 20180409. doi: 10.1371/journal.ppat.1006960. PubMed PMID: 29630679; PMCID: PMC5908200.

10. Weiss GE, Traore B, Kayentao K, Ongoiba A, Doumbo S, Doumtabe D, Kone Y, Dia S, Guindo A, Traore A, Huang CY, Miura K, Mircetic M, Li S, Baughman A, Narum DL, Miller LH, Doumbo OK, Pierce SK, Crompton PD. The Plasmodium falciparum-specific human memory B cell compartment expands gradually with repeated malaria infections. PLoS Pathog. 2010;6(5):e1000912. Epub 20100520. doi: 10.1371/journal.ppat.1000912. PubMed PMID: 20502681; PMCID: PMC2873912.

11. Nogaro SI, Hafalla JC, Walther B, Remarque EJ, Tetteh KK, Conway DJ, Riley EM, Walther M. The breadth, but not the magnitude, of circulating memory B cell responses to P. falciparum increases with age/exposure in an area of low transmission. PLoS One. 2011;6(10):e25582. Epub 20111004. doi: 10.1371/journal.pone.0025582. PubMed PMID: 21991321; PMCID: PMC3186790.

12. Wendel BS, He C, Qu M, Wu D, Hernandez SM, Ma KY, Liu EW, Xiao J, Crompton PD, Pierce SK, Ren P, Chen K, Jiang N. Accurate immune repertoire sequencing reveals malaria infection driven antibody lineage diversification in young children. Nat Commun. 2017;8(1):531. Epub 20170914. doi: 10.1038/s41467-017-00645-x. PubMed PMID: 28912592; PMCID: PMC5599618.

13. Marsh K, Otoo L, Hayes RJ, Carson DC, Greenwood BM. Antibodies to blood stage antigens of Plasmodium falciparum in rural Gambians and their relation to protection against infection. Trans R Soc Trop Med Hyg. 1989;83(3):293–303. doi: 10.1016/0035-9203(89)90478-1. PubMed PMID: 2694458.

14. Tan J, Piccoli L, Lanzavecchia A. The Antibody Response to Plasmodium falciparum: Cues for Vaccine Design and the Discovery of Receptor-Based Antibodies. Annu Rev Immunol. 2019;37:225–46. Epub 20181219. doi: 10.1146/annurev-immunol-042617-053301. PubMed PMID: 30566366.

15. Wu L, Mwesigwa J, Affara M, Bah M, Correa S, Hall T, Singh SK, Beeson JG, Tetteh KKA, Kleinschmidt I, D’Alessandro U, Drakeley C. Antibody responses to a suite of novel serological markers for malaria surveillance demonstrate strong correlation with clinical and parasitological infection across seasons and transmission settings in The Gambia. BMC Med. 2020;18(1):304. Epub 20200925. doi: 10.1186/s12916-020-01724-5. PubMed PMID: 32972398; PMCID: PMC7517687.

16. Kurup SP, Obeng-Adjei N, Anthony SM, Traore B, Doumbo OK, Butler NS, Crompton PD, Harty JT. Regulatory T cells impede acute and long-term immunity to blood-stage malaria through CTLA-4. Nat Med. 2017;23(10):1220–5. Epub 20170911. doi: 10.1038/nm.4395. PubMed PMID: 28892065; PMCID: PMC5649372.

17. Illingworth J, Butler NS, Roetynck S, Mwacharo J, Pierce SK, Bejon P, Crompton PD, Marsh K, Ndungu FM. Chronic exposure to Plasmodium falciparum is associated with phenotypic evidence of B and T cell exhaustion. J Immunol. 2013;190(3):1038–47. Epub 20121221. doi: 10.4049/jimmunol.1202438. PubMed PMID: 23264654; PMCID: PMC3549224.

18. Weiss GE, Crompton PD, Li S, Walsh LA, Moir S, Traore B, Kayentao K, Ongoiba A, Doumbo OK, Pierce SK. Atypical memory B cells are greatly expanded in individuals living in a malaria-endemic area. J Immunol. 2009;183(3):2176–82. Epub 20090710. doi: 10.4049/jimmunol.0901297. PubMed PMID: 19592645; PMCID: PMC2713793.

19. Scholzen A, Sauerwein RW. How malaria modulates memory: Activation and dysregulation of B cells in Plasmodium infection. Trends in Parasitology: Elsevier Ltd; 2013. p. 252–62.

20. Portugal S, Tipton CM, Sohn H, Kone Y, Wang J, Li S, Skinner J, Virtaneva K, Sturdevant DE, Porcella SF, Doumbo OK, Doumbo S, Kayentao K, Ongoiba A, Traore B, Sanz I, Pierce SK, Crompton PD. Malaria-associated atypical memory B cells exhibit markedly reduced B cell receptor signaling and effector function. Elife. 2015;4. Epub 20150508. doi: 10.7554/eLife.07218. PubMed PMID: 25955968; PMCID: PMC4444601.

21. Crompton PD, Kayala MA, Traore B, Kayentao K, Ongoiba A, Weiss GE, Molina DM, Burk CR, Waisberg M, Jasinskas A, Tan X, Doumbo S, Doumtabe D, Kone Y, Narum DL, Liang X, Doumbo OK, Miller LH, Doolan DL, Baldi P, Felgner PL, Pierce SK. A prospective analysis of the Ab response to Plasmodium falciparum before and after a malaria season by protein microarray. Proc Natl Acad Sci U S A. 2010;107(15):6958–63. Epub 20100329. doi: 10.1073/pnas.1001323107. PubMed PMID: 20351286; PMCID: PMC2872454.

22. Baird JK, Basri H, Weina P, MaGuire JD, Barcus MJ, Picarema H, Elyazar IR, Ayomi E, Sekartuti. Adult Javanese migrants to Indonesian Papua at high risk of severe disease caused by malaria. Epidemiol Infect. 2003;131(1):791–7. doi: 10.1017/s0950268803008422. PubMed PMID: 12948380; PMCID: PMC2870021.

23. Baird JK, Krisin, Barcus MJ, Elyazar IR, Bangs MJ, Maguire JD, Fryauff DJ, Richie TL, Sekartuti, Kalalo W. Onset of clinical immunity to Plasmodium falciparum among Javanese migrants to Indonesian Papua. Ann Trop Med Parasitol. 2003;97(6):557–64. doi: 10.1179/000349803225001472. PubMed PMID: 14511553.

24. Supargiyono S, Bretscher MT, Wijayanti MA, Sutanto I, Nugraheni D, Rozqie R, Kosasih AA, Sulistyawati S, Hawley WA, Lobo NF, Cook J, Drakeley CJ. Seasonal changes in the antibody responses against Plasmodium falciparum merozoite surface antigens in areas of differing malaria endemicity in Indonesia. Malar J. 2013;12:444. Epub 20131209. doi: 10.1186/1475-2875-12-444. PubMed PMID: 24321092; PMCID: PMC3866602.

25. Bediako Y, Ngoi JM, Nyangweso G, Wambua J, Opiyo M, Nduati EW, Bejon P, Marsh K, Ndungu FM. The effect of declining exposure on T cell-mediated immunity to Plasmodium falciparum - an epidemiological “natural experiment”. BMC Med. 2016;14(1):143. Epub 20160922. doi: 10.1186/s12916-016-0683-6. PubMed PMID: 27660116; PMCID: PMC5034532.

26. Tran TM, Jones MB, Ongoiba A, Bijker EM, Schats R, Venepally P, Skinner J, Doumbo S, Quinten E, Visser LG, Whalen E, Presnell S, O’Connell EM, Kayentao K, Doumbo OK, Chaussabel D, Lorenzi H, Nutman TB, Ottenhoff TH, Haks MC, Traore B, Kirkness EF, Sauerwein RW, Crompton PD. Transcriptomic evidence for modulation of host inflammatory responses during febrile Plasmodium falciparum malaria. Sci Rep. 2016;6:31291. Epub 20160810. doi: 10.1038/srep31291. PubMed PMID: 27506615; PMCID: PMC4978957.

27. Bediako Y, Adams R, Reid AJ, Valletta JJ, Ndungu FM, Sodenkamp J, Mwacharo J, Ngoi JM, Kimani D, Kai O, Wambua J, Nyangweso G, de Villiers EP, Sanders M, Lotkowska ME, Lin JW, Manni S, Addy JWG, Recker M, Newbold C, Berriman M, Bejon P, Marsh K, Langhorne J. Repeated clinical malaria episodes are associated with modification of the immune system in children. BMC Med. 2019;17(1):60. Epub 20190313. doi: 10.1186/s12916-019-1292-y. PubMed PMID: 30862316; PMCID: PMC6415347.

28. Warimwe GM, Keane TM, Fegan G, Musyoki JN, Newton CR, Pain A, Berriman M, Marsh K, Bull PC. Plasmodium falciparum var gene expression is modified by host immunity. Proc Natl Acad Sci U S A. 2009;106(51):21801–6. Epub 20091211. doi: 10.1073/pnas.0907590106. PubMed PMID: 20018734; PMCID: PMC2792160.

29. Schneider VM, Visone JE, Harris CT, Florini F, Hadjimichael E, Zhang X, Gross MR, Rhee KY, Ben Mamoun C, Kafsack BFC, Deitsch KW. The human malaria parasite Plasmodium falciparum can sense environmental changes and respond by antigenic switching. Proc Natl Acad Sci U S A. 2023;120(17):e2302152120. Epub 20230417. doi: 10.1073/pnas.2302152120. PubMed PMID: 37068249; PMCID: PMC10151525.

30. Coulibaly D, Travassos MA, Kone AK, Tolo Y, Laurens MB, Traore K, Diarra I, Niangaly A, Daou M, Dembele A, Sissoko M, Guindo B, Douyon R, Guindo A, Kouriba B, Sissoko MS, Sagara I, Plowe CV, Doumbo OK, Thera MA. Stable malaria incidence despite scaling up control strategies in a malaria vaccine-testing site in Mali. Malaria Journal. 2014;13:1–9. doi: 10.1186/1475-2875-13-374. PubMed PMID: 25238721.

31. Schall TJ, Jognstra J, Dyer BJ, Jorgensen J, Clayberger C, Davis MM, Krensky AM. A human T cell-specific molecule is a member of a new gene family. Journal of Immunology. 1988;141(3):1018–25.

32. Martinez-Navio JM, Casanova V, Pacheco R, Naval-Macabuhay I, Climent N, Garcia F, Gatell JM, Mallol J, Gallart T, Lluis C, Franco R. Adenosine deaminase potentiates the generation of effector, memory, and regulatory CD4+ T cells. J Leukoc Biol. 2011;89(1):127–36. Epub 20101019. doi: 10.1189/jlb.1009696. PubMed PMID: 20959412.

33. Pena SV, Krensky AM. Granulysin, a new human cytolytic granule-associated protein with possible involvement in cell-mediated cytotoxicity. Semin Immunol. 1997;9(2):117–25. doi: 10.1006/smim.1997.0061. PubMed PMID: 9194222.

34. Kuepper M, Koester K, Bratke K, Myrtek D, Ogawa K, Nagata K, Virchow JC, Jr., Luttmann W. Increase in Ksp37-positive peripheral blood lymphocytes in mild extrinsic asthma. Clin Exp Immunol. 2004;137(2):359–65. doi: 10.1111/j.1365-2249.2004.02540.x. PubMed PMID: 15270853; PMCID: PMC1809105.

35. Ihim SA, Abubakar SD, Zian Z, Sasaki T, Saffarioun M, Maleknia S, Azizi G. Interleukin- 18 cytokine in immunity, inflammation, and autoimmunity: Biological role in induction, regulation, and treatment. Front Immunol. 2022;13:919973. Epub 20220811. doi: 10.3389/fimmu.2022.919973. PubMed PMID: 36032110; PMCID: PMC9410767.

36. Haque M, Siegel RJ, Fox DA, Ahmed S. Interferon-stimulated GTPases in autoimmune and inflammatory diseases: promising role for the guanylate-binding protein (GBP) family. Rheumatology (Oxford). 2021;60(2):494–506. doi: 10.1093/rheumatology/keaa609. PubMed PMID: 33159795; PMCID: PMC7850581.

37. Iwata H, Goettsch C, Sharma A, Ricchiuto P, Goh WW, Halu A, Yamada I, Yoshida H, Hara T, Wei M, Inoue N, Fukuda D, Mojcher A, Mattson PC, Barabasi AL, Boothby M, Aikawa E, Singh SA, Aikawa M. PARP9 and PARP14 cross-regulate macrophage activation via STAT1 ADP-ribosylation. Nat Commun. 2016;7:12849. Epub 20161031. doi: 10.1038/ncomms12849. PubMed PMID: 27796300; PMCID: PMC5095532.

38. He G, Ma Y, Chou SY, Li H, Yang C, Chuang JZ, Sung CH, Ding A. Role of CLIC4 in the host innate responses to bacterial lipopolysaccharide. Eur J Immunol. 2011;41(5):1221–30. Epub 20110420. doi: 10.1002/eji.201041266. PubMed PMID: 21469130; PMCID: PMC3099427.

39. Yan R, Liu Z. LRRK2 enhances Nod1/2-mediated inflammatory cytokine production by promoting Rip2 phosphorylation. Protein Cell. 2017;8(1):55–66. Epub 20161109. doi: 10.1007/s13238-016-0326-x. PubMed PMID: 27830463; PMCID: PMC5233611.

40. Newman AM, Steen CB, Liu CL, Gentles AJ, Chaudhuri AA, Scherer F, Khodadoust MS, Esfahani MS, Luca BA, Steiner D, Diehn M, Alizadeh AA. Determining cell type abundance and expression from bulk tissues with digital cytometry. Nat Biotechnol. 2019;37(7):773–82. Epub 20190506. doi: 10.1038/s41587-019-0114-2. PubMed PMID: 31061481; PMCID: PMC6610714.

41. Nakayama T, Hirahara K, Kimura MY, Iwamura C, Kiuchi M, Kokubo K, Onodera A, Hashimoto K, Motohashi S. CD4+ T cells in inflammatory diseases: pathogenic T-helper cells and the CD69-Myl9 system. Int Immunol. 2021;33(12):699–704. doi: 10.1093/intimm/dxab053. PubMed PMID: 34427648.

42. Colborn JM, Ylostalo JH, Koita OA, Cisse OH, Krogstad DJ. Human Gene Expression in Uncomplicated Plasmodium falciparum Malaria. J Immunol Res. 2015;2015:162639. Epub 20150930. doi: 10.1155/2015/162639. PubMed PMID: 26491700; PMCID: PMC4605373.

43. Maier AG, Rug M, O’Neill MT, Brown M, Chakravorty S, Szestak T, Chesson J, Wu Y, Hughes K, Coppel RL, Newbold C, Beeson JG, Craig A, Crabb BS, Cowman AF. Exported proteins required for virulence and rigidity of Plasmodium falciparum-infected human erythrocytes. Cell. 2008;134(1):48–61. doi: 10.1016/j.cell.2008.04.051. PubMed PMID: 18614010; PMCID: PMC2568870.

44. Carmo OMS, Shami GJ, Cox D, Liu B, Blanch AJ, Tiash S, Tilley L, Dixon MWA. Deletion of the Plasmodium falciparum exported protein PTP7 leads to Maurer’s clefts vesiculation, host cell remodeling defects, and loss of surface presentation of EMP1. PLoS Pathog. 2022;18(8):e1009882. Epub 20220805. doi: 10.1371/journal.ppat.1009882. PubMed PMID: 35930605; PMCID: PMC9385048.

45. Stortz JF, Del Rosario M, Singer M, Wilkes JM, Meissner M, Das S. Formin-2 drives polymerisation of actin filaments enabling segregation of apicoplasts and cytokinesis in Plasmodium falciparum. Elife. 2019;8. Epub 20190719. doi: 10.7554/eLife.49030. PubMed PMID: 31322501; PMCID: PMC6688858.

46. White M, Watson J. Age, exposure and immunity. Elife. 2018;7:1-3. Epub 20180821. doi: 10.7554/eLife.40150. PubMed PMID: 30129437; PMCID: PMC6103766.

47. Ryg-Cornejo V, Ly A, Hansen DS. Immunological processes underlying the slow acquisition of humoral immunity to malaria. Parasitology. 2016;143(2):199–207. Epub 20160108. doi: 10.1017/S0031182015001705. PubMed PMID: 26743747.

48. Hviid L. Naturally acquired immunity to Plasmodium falciparum malaria in Africa. Acta Tropica2005. p. 270–5.

49. Baird JK. Age-dependent characteristics of protection v. susceptibility to Plasmodium falciparum. Annals of Tropical Medicine & Parasitology. 1997;92(4):367–90.

50. Brillantes M, Beaulieu AM. Memory and Memory-Like NK Cell Responses to Microbial Pathogens. Front Cell Infect Microbiol. 2020;10:102. Epub 20200325. doi: 10.3389/fcimb.2020.00102. PubMed PMID: 32269968; PMCID: PMC7109401.

51. Hart GT, Tran TM, Theorell J, Schlums H, Arora G, Rajagopalan S, Jules Sangala AD, Welsh KJ, Traore B, Pierce SK, Crompton PD, Bryceson YT, Long EO. Adaptive NK cells in people exposed to Plasmodium falciparum correlate with protection from malaria. Journal of Experimental Medicine 2019. p. 1280–90.

52. Arora G, Hart GT, Manzella-Lapeira J, Doritchamou JY, Narum DL, Thomas LM, Brzostowski J, Rajagopalan S, Doumbo OK, Traore B, Miller LH, Pierce SK, Duffy PE, Crompton PD, Desai SA, Long EO. NK cells inhibit Plasmodium falciparum growth in red blood cells via antibody-dependent cellular cytotoxicity. Elife. 2018;7. Epub 20180626. doi: 10.7554/eLife.36806. PubMed PMID: 29943728; PMCID: PMC6019063.

53. van den Hoogen LL, Walk J, Oulton T, Reuling IJ, Reiling L, Beeson JG, Coppel RL, Singh SK, Draper SJ, Bousema T, Drakeley C, Sauerwein R, Tetteh KKA. Antibody Responses to Antigenic Targets of Recent Exposure Are Associated With Low-Density Parasitemia in Controlled Human Plasmodium falciparum Infections. Front Microbiol. 2018;9:3300. Epub 20190116. doi: 10.3389/fmicb.2018.03300. PubMed PMID: 30700984; PMCID: PMC6343524.

54. Richards JS, Stanisic DI, Fowkes FJ, Tavul L, Dabod E, Thompson JK, Kumar S, Chitnis CE, Narum DL, Michon P, Siba PM, Cowman AF, Mueller I, Beeson JG. Association between naturally acquired antibodies to erythrocyte-binding antigens of Plasmodium falciparum and protection from malaria and high-density parasitemia. Clin Infect Dis. 2010;51(8):e50–60. doi: 10.1086/656413. PubMed PMID: 20843207.

55. Holz LE, Fernandez-Ruiz D, Heath WR. Protective immunity to liver-stage malaria. Clin Transl Immunology. 2016;5(10):e105. Epub 20161021. doi: 10.1038/cti.2016.60. PubMed PMID: 27867517; PMCID: PMC5099428.

56. Sutton HJ, Aye R, Idris AH, Vistein R, Nduati E, Kai O, Mwacharo J, Li X, Gao X, Andrews TD, Koutsakos M, Nguyen THO, Nekrasov M, Milburn P, Eltahla A, Berry AA, Kc N, Chakravarty S, Sim BKL, Wheatley AK, Kent SJ, Hoffman SL, Lyke KE, Bejon P, Luciani F, Kedzierska K, Seder RA, Ndungu FM, Cockburn IA. Atypical B cells are part of an alternative lineage of B cells that participates in responses to vaccination and infection in humans. Cell Rep. 2021;34(6):108684. doi: 10.1016/j.celrep.2020.108684. PubMed PMID: 33567273; PMCID: PMC7873835.

57. Perez-Mazliah D, Gardner PJ, Schweighoffer E, McLaughlin S, Hosking C, Tumwine I, Davis RS, Potocnik AJ, Tybulewicz VL, Langhorne J. Plasmodium-specific atypical memory B cells are short-lived activated B cells. Elife. 2018;7. Epub 20181102. doi: 10.7554/eLife.39800. PubMed PMID: 30387712; PMCID: PMC6242553.

58. Rodriguez-Barraquer I, Arinaitwe E, Jagannathan P, Kamya MR, Rosenthal PJ, Rek J, Dorsey G, Nankabirwa J, Staedke SG, Kilama M, Drakeley C, Ssewanyana I, Smith DL, Greenhouse B. Quantification of anti-parasite and anti-disease immunity to malaria as a function of age and exposure. Elife. 2018;7. Epub 20180725. doi: 10.7554/eLife.35832. PubMed PMID: 30044224; PMCID: PMC6103767.

59. Rodriguez-Barraquer I, Arinaitwe E, Jagannathan P, Boyle MJ, Tappero J, Muhindo M, Kamya MR, Dorsey G, Drakeley C, Ssewanyana I, Smith DL, Greenhouse B. Quantifying Heterogeneous Malaria Exposure and Clinical Protection in a Cohort of Ugandan Children. J Infect Dis. 2016;214(7):1072–80. Epub 20160801. doi: 10.1093/infdis/jiw301. PubMed PMID: 27481862; PMCID: PMC5021229.

60. Andrade CM, Fleckenstein H, Thomson-Luque R, Doumbo S, Lima NF, Anderson C, Hibbert J, Hopp CS, Tran TM, Li S, Niangaly M, Cisse H, Doumtabe D, Skinner J, Sturdevant D, Ricklefs S, Virtaneva K, Asghar M, Homann MV, Turner L, Martins J, Allman EL, N’Dri ME, Winkler V, Llinas M, Lavazec C, Martens C, Farnert A, Kayentao K, Ongoiba A, Lavstsen T, Osorio NS, Otto TD, Recker M, Traore B, Crompton PD, Portugal S. Increased circulation time of Plasmodium falciparum underlies persistent asymptomatic infection in the dry season. Nat Med. 2020;26(12):1929–40. Epub 20201026. doi: 10.1038/s41591-020-1084-0. PubMed PMID: 33106664.

61. Baird JK. Host Age as a Determinant of Naturally Acquired Immunity to Plasmodium falciparum. Parasitology Today. 1995;11(3):105–11.

62. Quin JE, Bujila I, Cherif M, Sanou GS, Qu Y, Vafa Homann M, Rolicka A, Sirima SB, O’Connell MA, Lennartsson A, Troye-Blomberg M, Nebie I, Ostlund Farrants AK. Major transcriptional changes observed in the Fulani, an ethnic group less susceptible to malaria. Elife. 2017;6. Epub 20170919. doi: 10.7554/eLife.29156. PubMed PMID: 28923166; PMCID: PMC5629023.

63. Kim D, Paggi JM, Park C, Bennett C, Salzberg SL. Graph-based genome alignment and genotyping with HISAT2 and HISAT-genotype. Nat Biotechnol. 2019;37(8):907–15. Epub 20190802. doi: 10.1038/s41587-019-0201-4. PubMed PMID: 31375807; PMCID: PMC7605509.

64. Aurrecoechea C, Brestelli J, Brunk BP, Dommer J, Fischer S, Gajria B, Gao X, Gingle A, Grant G, Harb OS, Heiges M, Innamorato F, Iodice J, Kissinger JC, Kraemer E, Li W, Miller JA, Nayak V, Pennington C, Pinney DF, Roos DS, Ross C, Stoeckert CJ, Treatman C, Wang H. PlasmoDB: A functional genomic database for malaria parasites. Nucleic Acids Research. 2009;37:539–43. doi: 10.1093/nar/gkn814.

65. Liao Y, Smyth GK, Shi W. featureCounts: an efficient general purpose program for assigning sequence reads to genomic features. Bioinformatics. 2014;30(7):923–30. Epub 20131113. doi: 10.1093/bioinformatics/btt656. PubMed PMID: 24227677.

66. Robinson MD, McCarthy DJ, Smyth GK. edgeR: a Bioconductor package for differential expression analysis of digital gene expression data. Bioinformatics. 2010;26(1):139–40. Epub 20091111. doi: 10.1093/bioinformatics/btp616. PubMed PMID: 19910308; PMCID: PMC2796818.

67. Benjamini Y, Hochberg Y. Controlling the False Discovery Rate : A Practical and Powerful Approach to Multiple Testing Author ( s ): Yoav Benjamini and Yosef Hochberg Source : Journal of the Royal Statistical Society . Series B ( Methodological ), Vol . 57 , No . 1 ( 1995 ), Publi. Journal of the Royal Statistical Society. 1995;57:289–300.

68. Chen B, Khodadoust MS, Liu CL, Newman AM, Alizadeh AA. Profiling Tumor Infiltrating Immune Cells with CIBERSORT. Methods Mol Biol. 2018;1711:243–59. doi: 10.1007/978-1-4939-7493-1_12. PubMed PMID: 29344893; PMCID: PMC5895181.

69. Tebben K, Dia A, Serre D. Determination of the Stage Composition of Plasmodium Infections from Bulk Gene Expression Data. mSystems. 2022;7(4):e0025822. Epub 20220705. doi: 10.1128/msystems.00258-22. PubMed PMID: 35862820; PMCID: PMC9426464.

70. Li H, Handsaker B, Wysoker A, Fennell T, Ruan J, Homer N, Marth G, Abecasis G, Durbin R, Genome Project Data Processing S. The Sequence Alignment/Map format and SAMtools. Bioinformatics. 2009;25(16):2078–9. Epub 20090608. doi: 10.1093/bioinformatics/btp352. PubMed PMID: 19505943; PMCID: PMC2723002.

71. Chan ER, Menard D, David PH, Ratsimbasoa A, Kim S, Chim P, Do C, Witkowski B, Mercereau-Puijalon O, Zimmerman PA, Serre D. Whole Genome Sequencing of Field Isolates Provides Robust Characterization of Genetic Diversity in Plasmodium vivax. PLoS Neglected Tropical Diseases 2012.

